# Morphological and functional variability in central and subcentral motor cortex of the human brain

**DOI:** 10.1101/2020.03.17.995035

**Authors:** Nicole Eichert, Kate E. Watkins, Rogier B. Mars, Michael Petrides

## Abstract

There is a long-established link between anatomy and function in the somatomotor system in the mammalian cerebral cortex. The morphology of the central sulcus is predictive of the location of functional activation peaks relating to movement of different effectors in individuals. By contrast, morphological variation in the subcentral region and its relationship to function is, as yet, unknown. Investigating the subcentral region is particularly important in the context of speech, since control of the larynx during human speech production is related to activity in this region. Here, we examined the relationship between morphology in the central and subcentral region and the location of functional activity during movement of the hand, lips, tongue, and larynx at the individual participant level. We provide a systematic description of the sulcal patterns of the subcentral and adjacent opercular cortex, including the inter-individual variability in sulcal morphology. We show that, in the majority of participants, the anterior subcentral sulcus is not continuous, but consists of two distinct segments. A robust relationship between morphology of the central and subcentral sulcal segments and movement of different effectors is demonstrated. Inter-individual variability of underlying anatomy might thus explain previous inconsistent findings, in particular regarding the ventral larynx area in subcentral cortex. A surface registration based on sulcal labels indicated that such anatomical information can improve the alignment of functional data for group studies.

## Introduction

A fundamental challenge in neuroscience is to establish meaningful links between brain structure and function. One of the clearest cases of such a structure-to-function relationship is found in the somatomotor system in the central strip of the cerebral cortex (Jackson 1863; Fritsch and Hitzig 1870; Vogt and Vogt 1919; Penfield and Boldrey 1937). Different parts of the body are represented in an orderly and consistent fashion, following a somatotopic organization, the schematic visualization of which is known as Penfield’s ‘homunculus’.

In contrast to electrical brain mapping studies, neuroimaging studies typically report average activations from larger groups of participants. This approach demonstrates effects that are representative of the human brain in general, but obscures participant-specific features and inter-individual variability, limiting sensitivity and functional resolution (Bennett and Miller 2010; Nieto-Castañón and Fedorenko 2012; Woo et al. 2014). Consequently, a growing number of neuroimaging studies now tend to report activation patterns in individuals (Miller et al. 2002; Barch et al. 2013; Carey et al. 2017). A deeper understanding of structure-to-function relationships, however, requires examination of inter-individual functional variability together with anatomical variability.

Several studies have established a coupling between the sulcal/gyral brain anatomy and functional activation in individuals (e.g., Amiez et al. 2006, 2013; Derrfuss et al. 2012; Zlatkina et al. 2016; Bodin et al. 2018). A recent neuroimaging study demonstrated that the location of individual somatomotor activation peaks held consistent relationships between the effector being moved and the morphologically defined segments of the central sulcus (Germann et al. 2020). The inter-individual variability of the central sulcus segments, however, was not quantified and this investigation focused exclusively on the central sulcus.

Movement representation in the human brain is not limited to the central sulcus. The subcentral gyrus, in addition to the ventral portion of the central somatomotor strip, is involved in speech-related movements (Penfield and Boldrey 1937; Olthoff et al. 2008; Grabski et al. 2012; Bouchard et al. 2013; Eichert et al. 2020a). Voluntary control of laryngeal movements during vocalization evokes brain activity in two distinct regions in lateral motor cortex: a dorsal region close to the representation of the lips and a ventral region close to the lateral fissure (reviewed in: Belyk and Brown 2017; Eichert et al. 2020a). The representation of the larynx in the human brain, however, remains controversial. The inconsistency among reports in the literature might be explained by inter-individual variability in structure and function in this region. Thus, examining the functional anatomy of the speech motor system requires investigation both of the central sulcus and the subcentral gyral region at an individual participant level.

To the best of our knowledge, there has been no systematic examination of the morphological variability in the subcentral gyrus and the adjacent opercular cortex. There are two distinct sulci in the subcentral region: the anterior subcentral sulcus (*ascs*) and the posterior subcentral sulcus (*pscs*) (Ono et al. 1990; Petrides 2012, 2019). The variability of these two sulci in relation to the segments of the central sulcus across individuals, and the relationship of morphological variability to functional brain activity during movement, specifically vocalization, remain to be established.

Here, we assessed the structure-to-function relationship in the human motor system on an individual participant level by examining the morphology of the central sulcus, the subcentral gyrus and the adjacent central operculum. We performed sulcal labeling in surface space, rather than in volume space, which allows for better visualization and surface-based registration with respect to the 2D topology of the cerebral cortex (Fischl et al. 1999b, a; Ghosh et al. 2010).

To visualize and quantify inter-individual variability of the examined sulcal segments, we derived spatial probability maps in both standard surface and volume stereotaxic space. In a subset of participants, we acquired functional magnetic resonance imaging (fMRI) data to localize brain activity during movement of the hand, lips and tongue, as well as the larynx during vocalization (Eichert et al. 2020a). To investigate how this structure-to-function relationship can help to improve alignment of data, we registered the brains of all participants based on the sulcal surface labels. A decrease in distances across activation peaks after applying the registration would indicate that anatomical variability can explain some of the functional variability.

## Material and Methods

### Participants

Data from two groups of participants were used. In one study group, both structural and functional MRI data were acquired (*n* = 20), but in the other study group, only structural MRI data were available (*n* = 30).

Structural and functional MRI data from the performance of motor tasks were acquired from 20 participants (12 females, 18 – 40 years [27.4 ± 5.6, mean ± SD], 5 self-reported left-handers). All participants were self-reported native English speakers (two were raised bilingually from infancy and three were fluent in a second acquired language) and had no history or diagnosis of speech disorders. All had normal hearing, normal or corrected-to-normal vision, and no neurological impairments. The participants were part of a study that had been approved by the Central University Research Ethics Committee of the University of Oxford (CUREC, R55787/RE001) in accordance with the regulatory standards of the Code of Ethics of the World Medical Association (Declaration of Helsinki). All participants gave informed consent for their participation and were monetarily compensated.

In addition, we used cortical brain surface reconstructions from 30 participants provided by the Human Connectome Project (HCP), WU-Minn Consortium (Principal Investigators: David Van Essen and Kamil Ugurbil; 1U54MH091657) funded by the 16 NIH Institutes and Centers that support the NIH Blueprint for Neuroscience Research; and by the McDonnell Center for Systems Neuroscience at Washington University (Van Essen et al. 2013). The minimally pre-processed datasets of the first 31 participants (16 female, age range 22-35 years, handedness information not accessed) of the Q2 release were used. One participant was excluded because of a technical problem in the automatic FreeSurfer parcellation.

### MRI Data Acquisition

MRI data acquisition parameters differed for the two groups of participants. Data from the participants that took part in the functional study were obtained at the Oxford Centre for Human Brain Activity (OHBA) using a 3T Siemens Prisma scanner with a 32-channel head coil. Two structural images of the whole brain had been acquired at 1 mm isotropic resolution: a T1w image (MPRAGE sequence) and a T2w image (SPACE sequence). For task-fMRI, whole head T2*-weighted echo planar images were acquired at 2.4 mm^3^ isotropic resolution (TE = 30 ms, multiband fact 6, TR = 0.8 s, Casey et al. 2018).

Details of data acquisition and preprocessing methods of the HCP participants are provided in Glasser et al. (2013) and Uǧurbil et al. (2013). T1w images had been acquired using an MPRAGE sequence at 0.7 mm isotropic resolution.

### Structural MRI Data Analysis

Data from the participants who took part in the functional study were pre-processed using the HCP-pipeline (Glasser et al. 2013). The automatic processing pipeline includes cortical surface reconstruction using FreeSurfer based on the contrast from the T1w and the T2w images and automatic assignment of neuroanatomical labels. Cortical surface reconstructions of the HCP participants were derived using FreeSurfer based on the T1w scans and directly provided by the database. A linear transformation (12 degrees of freedom) from the high-resolution T1w anatomical scans to standard MNI space (nonlinear 6^th^ generation atlas, Fonov et al. 2011) was derived using FSL’s FLIRT (Jenkinson and Smith 2001; Jenkinson et al. 2002) and further refined using FNIRT nonlinear registration (Andersson et al. 2007).

### Sulcal Labeling

We identified the following sulci and sulcal segments in the structural data from all 50 participants: five segments of the central sulcus from dorsal to ventral (*cs_1* to *cs_5*), the lateral and opercular segments of the anterior subcentral sulcus (*ascs_lat, ascs_op*) and the posterior subcentral sulcus (*pscs*) (Fig. 1a). Sulcal labels for one example participant are shown in Fig. 1b and Fig. 1c and a three-dimensional rendering of this individual’s segments is provided as animation in the supplementary material (Online Resource 1) generated using FSLeyes (McCarthy 2020). Note that all sulcal labels and surface reconstructions will be made openly accessible to allow interactive inspection in a 3D viewer. Sulcal labels were drawn manually onto the native surface mesh (approximately 136,000 vertices) in Connectome Workbench’s wb_view (www.humanconnectome.org/software/connectome-workbench.html). Surface features of both pial and white matter surfaces were inspected in conjunction with the participant’s T1w scan. The identification of the central sulcus segments was based on changes in direction of the sulcus, and on the location of gyral ‘plis de passage’, which are small gyral bridges connecting the postcentral with the precentral gyrus, or based on more subtle ‘wall pinches’. These landmarks can be most easily identified on the white matter surface and are not always visible on the pial surface alone (Germann et al. 2020). The following description of morphological criteria was sufficient to identify the sulci in all 50 participants.

**Fig. 1.**
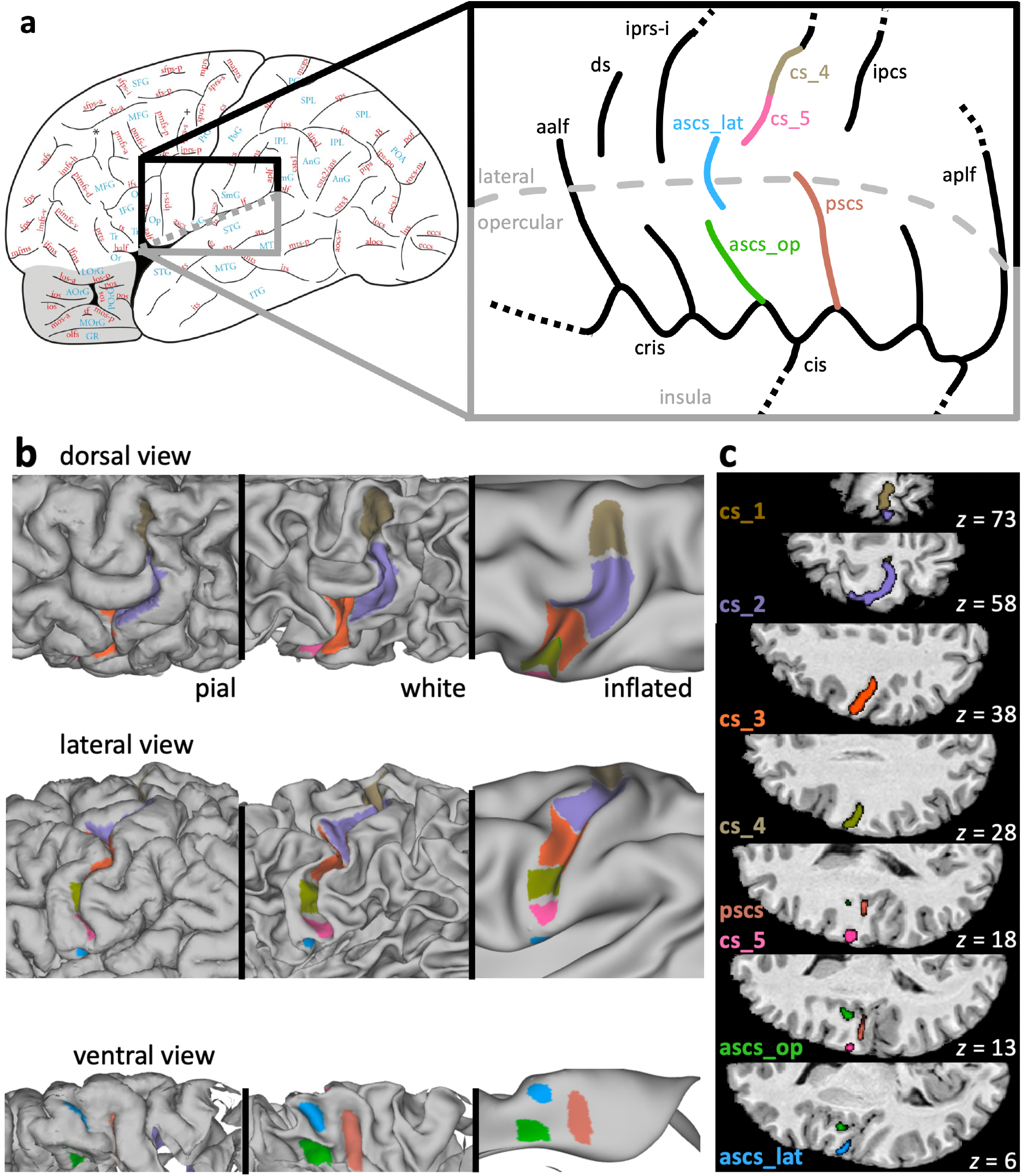
Sulcal morphology in the central and the subcentral region. **a** Left: Sulcal map of the human cerebral cortex (reprinted from Petrides M. (2019) Atlas of the morphology of the human cerebral cortex on the average MNI brain. New York: Academic Press, with permission from Elsevier). The region shown in grey depicts the orbitofrontal cortex. Right: Typical configuration of sulci in the subcentral region. This configuration was classified as morphological Type 1 (see Fig. 2). Anatomical variability was assessed in the sulci marked with color: *ascs_lat*, lateral segment of the anterior subcentral sulcus; *ascs_op*, opercular segment of the anterior subcentral sulcus; *cs_1* to *cs_5*, segments of the central sulcus (only *cs_4* and *cs_5* are shown in the highlighted region); *pscs*, posterior subcentral sulcus. Grey dashed line: visible surface boundary between lateral and opercular cortex. *aalf*, ascending anterior ramus of the lateral fissure; *aplf*, ascending posterior ramus of the lateral fissure; *cis*, central insular sulcus; *cris*, circular insular sulcus; *ds*, diagonal sulcus; *iprs*, inferior precentral sulcus. **b** Sulcal labels in one individual displayed onto the native pial surface, white matter surface, and inflated surface. **c** Sulcal labels of the same participant in MNI volume space

*Cs_1* is the most dorsal segment of the central sulcus, which runs more or less in a vertical straight direction. Its ventral boundary was drawn at the location where a gyral bridge provides a prominent landmark on the posterior bank of the central sulcus. *Cs_2* has a characteristic curvature in the shape of the Greek omega letter (see Fig. 1b, c), which is known as the ‘hand knob’ (Yousry et al. 1997). The hand knob can be split into two smaller knob-like curves, the so-called epsilon configuration (Yousry et al. 1997). If this configuration were present, both knobs were labeled together as *cs_2* and the pinch on the posterior wall of the central sulcus in the middle was not treated as a boundary between these labels. The boundary between *cs_2* and *cs_3* was drawn at the location where the central sulcus changes direction and where a gyral passage can be observed on the posterior bank. In some brains, an additional convexity of the central sulcus can be observed on the posterior bank in the middle of *cs_3*. The ventral boundary of *cs_3* was drawn ventral to this convexity, if present, at the location where a small gyral bridge forms a landmark on the anterior bank of the central sulcus. The last two segments of the central sulcus, *cs_4* and *cs_5* are smaller in extent, shallower and more variable in their morphology (see Results section for a more detailed anatomical characterization). The boundary between *cs_4* and *cs_5* was defined based on a gyral bridge on the posterior bank of the central sulcus. *Cs_5*, which is the most ventral part of the central sulcus, can form an additional curve or run straight in a variable direction.

The labels for *ascs* and *pscs* were assigned based on an atlas of human brain morphology (Petrides, 2019). For *ascs*, we labeled two distinct segments: a lateral and an opercular segment (*ascs_lat, ascs_op*). The course of *ascs_lat*, *ascs_op* and *pscs* was highly variable and a detailed description of the sulcal anatomy in the subcentral region is reported in the results section.

The morphological patterns of the ventral subcentral region were categorized into five types depending on the configuration of the *ascs_lat*. The classification was based on the location of the *ascs_lat* on the cortex and its spatial relation to other sulci. Sulcal segments were considered as ‘merged’, when there was a clear continuation on the pial surface, although, in some cases, a discontinuity between the merged sulci was still observed on the white matter surface.

### Spatial Probability Maps

In order to characterize the inter-individual morphological variability of the labeled sulci, we generated probability maps in surface and volume space. To obtain surface probability maps, all surface labels were resampled from native to a standard mesh (32k_fs_LR) following FreeSurfer’s registration (Glasser et al. 2013). At each vertex, the 50 surface maps were binarized, summed up and then normalized to create a surface label with intensities ranging from 0 % to 100 % at the maximal possible overlap of all 50 participants. For visualization, the surface probability maps were displayed onto an inflated template surface.

For the generation of volumetric probability maps, individual surface labels were mapped to volume space. For surface-to-volume mapping, we used the participant’s native pial surface registered to 32k_fs_LR mesh and non-linearly transformed to MNI-space (i.e. the surfaces in the subfolder /MNINonLinear/fsaverage_LR32k/). We performed the mapping to 0.5 mm resolution volume space using wb_command and ‘-nearest-vertex’ option. Given that the volumetric labels spatially varied in 3 rather than in 2 dimensions, the labels were smoothed (Gaussian kernel with FWHM of 2 mm) to enhance overlap. The smoothed labels were then thresholded at 0.1, binarized and then summed at each voxel. For visualization, volume probability maps were overlaid onto the MNI average brain.

### Anatomical characterization of sulcal segments

We characterized the sulcal segments using several anatomical descriptors. Inter-individual variability was determined as the maximal value within the surface probability map. For the measure of sulcal depth, we computed the minimal value in each individual’s sulcal segment labels using FreeSurfer’s sulcal depth map. The depth value is based on an individual’s pial surface and expressed as normalized value relative to a baseline running along the sulcal banks, rather than in real-world units. The depth values range from positive values at the crown of a gyrus to negative values at the fundus of a sulcus. Furthermore, we computed the mean cortical thickness in each label, as provided by FreeSurfer. As measure for the size of the sulcal segment, we computed the number of vertices that the surface label spanned on the native surface.

### Functional MRI Experimental Design

The 20 participants who provided structural and functional data took part in an fMRI study on speech production and laryngeal motor control. The experimental design, processing, and fMRI results of this study have been reported elsewhere in detail (Eichert et al. 2020a) and are here only briefly described. In a functional localizer task, participants were asked to perform repeated lip protrusion or tongue retraction at a rate of approximately 1-2 reps/s. The participant’s breathing pattern was explicitly controlled using the fixation symbol on the screen, instructing them to inhale for 1.5 s and exhale for 4 s. A ‘breathing only’ condition, during which the participants followed the same breathing pattern, was acquired as baseline condition. Each task condition was performed in blocks lasting 22 s followed by a rest period of 8 s with normal breathing. The conditions were presented in a fixed pseudo-random order following a balanced Latin-square design wherein each condition was repeated four times.

In a separate task, participants were instructed to produce a syllable sequence (/la leɪ li la leɪ li/) under four different conditions: overt speech, silent mouthing, only vowel production, and covert speech. Breathing instructions, task timing and randomization of the four blocks were the same as described for the localizer task, except that each condition was repeated five times. In a third task, participants performed a task that required both phonological and semantic judgements. Participants had to indicate a yes/no response by pressing a button with the right index or the middle finger every 3 s. This task was analyzed as a localizer for the hand region in the left hemisphere.

### Functional MRI Data Analysis and Statistical Analysis

Functional MRI data processing was carried out using FEAT (FMRI Expert Analysis Tool) Version 6.00, part of FSL (FMRIB’s Software Library, www.fmrib.ox.ac.uk/fsl) including motion correction of the images and unwarping using a fieldmap (Jenkinson 2003). Time-series statistical analysis was based on a general linear model (GLM) implemented in FILM with local autocorrelation correction (Woolrich et al. 2001). Standard motion correction parameters and individual volumes that were motion outliers, determined using fsl_motion_outliers, were included as separate regressors at the first level for each participant. Registration to the high-resolution structural scan and standard 2-mm MNI template was carried out using FLIRT. Registration from high resolution structural to MNI space was then further refined using FNIRT nonlinear registration (Andersson et al. 2007).

In the functional localizer task for lip and tongue movements, activity during each condition was assessed relative to the ‘breathing only’ condition. For the syllable production task, the conditions were analyzed in a factorial model that allowed separation of the (supra-laryngeal) articulation and the (laryngeal) vocalization components of the task. Brain activity associated with the control of supra-laryngeal articulation was defined as (‘overt speech’ *minus* ‘vowel production’) *plus* (‘silent mouthing’ *minus* ‘covert speech’) and the main contrast for vocalization was derived from the contrast (‘overt speech’ *minus* ‘silent mouthing’) *plus* (‘vowel production’ *minus* ‘covert speech’). The task activations from the ‘articulation’ contrast are not further discussed in this manuscript.

In both tasks described above, the rest blocks with normal breathing served as baseline, which means that they were not modelled in the GLM. For the hand localizer task, we derived a contrast of all conditions involving button presses relative to a resting baseline. Note that this task only provided data to analyze the hand representation in the left hemisphere.

### Individual Surface Activation Maxima

To assess inter-individual variability of the fMRI results, we derived the location of individual activation maxima for hand, lip, and tongue movements, and larynx activity during vocalization. Activation maxima were derived using the steps described in Eichert et al. (2020a), and are reported here only briefly. ROI definitions are described in more detail in the supplementary material.

Different volumetric ROI masks were used for the different motor representations based on individual anatomy in both hemispheres. We used an ROI of the whole central sulcus for hand, lip and tongue, based on the Destrieux Atlas. For the dorsal larynx representation, we used a more limited portion of the central sulcus ROI (MNI *z*-coordinates: 50 – 30). For the ventral larynx representation, we manually defined an ROI based on individual surface landmarks.

Individual volumetric ROIs were linearly transformed from FreeSurfer’s anatomical to functional space of the respective task fMRI scan. Within the ROI, the voxel of maximal intensity was determined from the uncorrected *z*-statistics image. Activation maxima were manually inspected in the participant’s native volume space to confirm that the systematic approach described below captured task-related activations. It should be noted that, for some participants, this local maximum did not achieve the corrected voxel-wise significance threshold (left hemisphere: hand *n* = 3, dorsal larynx *n* = 6, ventral larynx *n* = 5; right hemisphere: dorsal larynx *n* = 5, ventral larynx *n* = 4). Using a lower uncorrected threshold is justified given our goal to visualize and assess spatial variability of the activation maxima. The activation maxima were mapped to the individual’s native mid-thickness surface, resampled to the 32k standard (fs_LR) surface mesh using the FreeSurfer registration, smoothed (FWHM = 1 mm), and binarized to form a small circular patch.

### Structure-to-Function Relationship

Next, we examined the spatial relationship between the sulcal segments, and the task activation peaks at the individual participant level. This analysis was performed in the 20 participants who contributed both task activation and structural data. Individual task activation peaks were mapped onto the individual’s cortical surface as described above for sulcal segments. In order to characterize the structure-to-function relationship at the group level, we aligned all individual surfaces based on the anatomical surface labels and then we applied the same registration to individual task activation peaks. This approach allowed us to visualize individual variability in the spatial distribution of task activation peaks in a common group-level space.

The registration of sulcal labels was driven by the binary labels for *cs_1*, *cs_2*, *cs_3*, *cs_4*, *cs_5*, *ascs_lat, ascs_op* and *pscs* and performed using the multimodal surface matching tool (MSM, Robinson et al. 2014), which is part of FSL. As target, or reference, for the MSM-based registration, we used the normalized and thresholded (> 0.4) average labels after projecting all of them to the same regular sphere (approximately 32,000 vertices) without using any other anatomical priors. Each participant’s sulcal maps and the reference sulcal maps were merged into a combined file with six data arrays, i.e. they were provided as metric ‘func.gii’ file with six maps. Then, we derived a registration using MSM for each participant. We used default settings for MSM and the following configuration parameters were empirically determined: levels=3; sigma_in=10,5,2; sigma_ref=10,5,2; lambda=0.1,0.01,0.01; it=10,10,3; opt=DISCRETE,DISCRETE,DISCRETE; CPgrid=2,3,4; SGgrid=4,5,6; datagrid=4,5,6.

We explored how manual labeling of sulcal segments can help to align data across participants. Thus, we compared our registration based on sulcal segments with the FreeSurfer registration and a registration based on sulcal depth maps. The FreeSurfer registration and sulcal depth maps were provided by the HCP-processing pipeline. To derive a registration based on sulcal depth, we used MSM default settings. To characterize the effect of the registrations, we quantified the spatial spread of the fMRI activation peaks: For each effector, we computed the median geodesic distances across all 20 activation peaks. The median distance for each effector was, therefore, based on 190 datapoints. A smaller spatial spread of activation peaks indicates a better registration performance.

## Results

### Sulcal Patterns in the Subcentral Region

The sulcal anatomy of the subcentral region was examined in a total of 100 hemispheres (*n* = 50 brains) and the morphological patterns encountered were classified into five types (Fig. 2). The most commonly observed configuration of subcentral sulci (Type 1, 44 % of hemispheres) is shown in Fig. 1b, c. In the Type 1 configuration, the central sulcus does not extend ventrally to meet the lateral fissure. The fifth and most ventral segment of the central sulcus (*cs_5*) is typically short and forms a curve in an anterior direction. The lateral segment of *ascs* (*ascs_lat*) is visible on the lateral surface of the brain and is clearly separate from the central sulcus. Dorsally, the *ascs_lat* extends into the precentral gyrus with variable length. Ventrally, the *ascs_lat* continues into the opercular cortex, which is hidden within the lateral fissure of the brain. A small gyral bridge separates the *ascs_lat* from the opercular segment of the *ascs* (*ascs_op*) which continues medially until it reaches the circular insular sulcus (*cris*). The medial origin of the *ascs_op* can be identified at a curve of *cris*, which is formed by the posterior short insular gyrus just anterior to the central insular sulcus (*cis*). Posterior to the *ascs_op*, the posterior subcentral gyrus (*pscs*) can be found, which extends laterally towards the lateral fissure. The medial origin of *pscs* can be identified at a curve of *cris* posterior to *cis*, which is formed by the anterior long insular gyrus. The number of curvatures of *cris* and the number of subcentral sulci is variable, but the *ascs_op* and the *pscs* could be reliably identified in every participant examined. The morphology of all of the mentioned sulci is variable across participants, but the description of morphological types reported below focusses on the configuration of the *ascs_lat*.

**Fig. 2.**
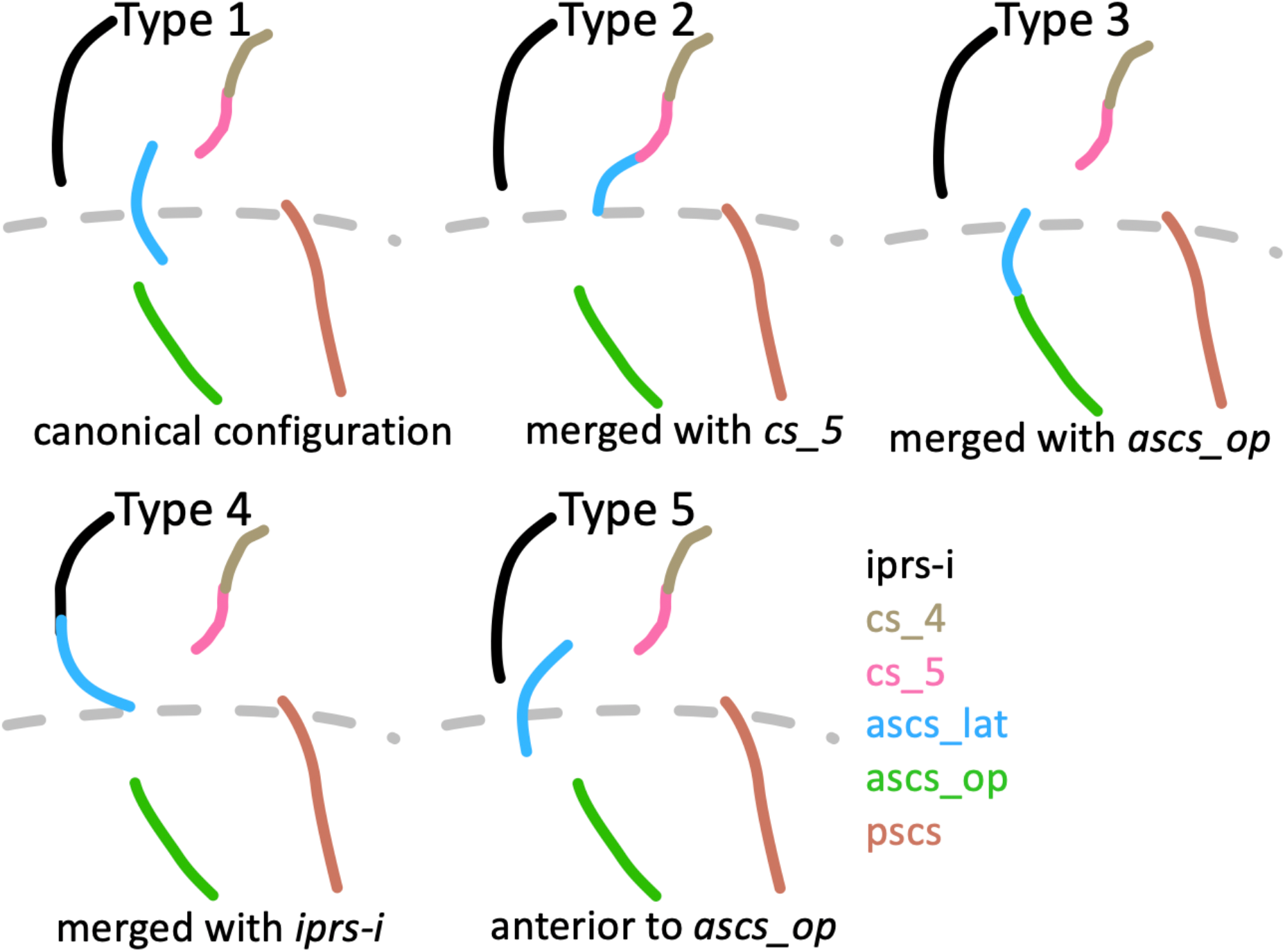
Morphological patterns. Schematic drawings illustrating the main morphological patterns of the subcentral region formed by the *ascs_lat* with neighbouring sulci. Type 1: Canonical configuration observed in the majority of hemispheres (also shown in Fig. 1b). Type 2: *ascs_lat* merging with the central sulcus. Type 3: *ascs_lat* merging with *ascs_op*. Type 4: *ascs_lat* merging with the inferior ramus of *iprs*. Type 5: *ascs_lat* anterior to *ascs_op*

### Morphological Patterns in the Subcentral Region

The subcentral region exhibits high inter-individual variability regarding the configuration of the *ascs_lat*. The observed occurrences of morphological types are reported in Table 1. In addition to the canonical (Type 1, 44 % of hemispheres) configuration described above, it was commonly observed that the *ascs_lat* merged with the central sulcus (Type 2, 20 %) (Fig. 2). In these cases, the *ascs_lat* remains mostly on the lateral brain surface and does not reach into the opercular cortex. The *ascs_op* extends further lateral and its extension is visible on the lateral brain surface. Another commonly observed type was identified by a more opercular position of the *ascs_lat* (Type 3, 18 %). In Type 3, the *ascs_lat* is less visible on the lateral surface and the central sulcus extends further ventral towards the lateral fissure. Type 3 is also characterized by a reduced gyral bridge separating the *ascs_lat* from the *ascs_op*. In six cases, the opercular continuation of the *ascs_lat* curved posteriorly and merged with the *pscs* instead of the *ascs_op* (not classified as a separate type). In several cases, we also observed the *ascs_lat* to merge with the inferior ramus of the inferior precentral sulcus (*iprs-i*) (Type 4). When the *ascs_lat* and the *iprs-i* merge, the ventral continuation of the *ascs_lat* curves in an anterior rather than a posterior direction and the position of the *ascs_lat* is more lateral than opercular. In a few cases (Type 5), the position of the *ascs_lat* was notably further rostral so that it was positioned anterior to the *ascs_op* rather than posterior as in the other configurations.

Strong hemispheric differences were observed in the occurrences of the morphological types (Table 1). Type 2 (*ascs_lat* mering with *cs_5*) is much more common in left hemispheres (85 % vs. 15 %), while Type 4 (*ascs_lat* merging with *iprs*) and Type 5 (*ascs_lat* further anterior), are more common in right hemispheres (75 % vs. 25 % for Type 4; 83 % vs. 17 % for Type 5), although both types are infrequent. For Type 1 (canonical configuration) and Type 3 (*ascs_lat* merging with *ascs_op*), no pronounced hemispheric differences were observed.

**Table 1.** Morphological types in the subcentral region. Observed frequencies of morphological patterns based on 100 examined hemispheres (LH – left hemisphere; RH – right hemisphere).

|  | Type 1 | Type 2 | Type 3 | Type 4 | Type 5 |
| --- | --- | --- | --- | --- | --- |
| LH | 20 | 17 | 9 | 3 | 1 |
| RH | 24 | 3 | 9 | 9 | 5 |
| Total | 44 | 20 | 18 | 12 | 6 |

### Spatial Probability Maps

The morphological variability of the central sulcus segments and the subcentral sulci was quantified and visualized using spatial probability maps in 2D surface space and 3D volumetric space (Fig. 3). For both surface and volume probability maps, intensity values decrease from the center of the map towards the edges, which is typical for overlap maps. Values in the volumetric probability maps are overall lower given that they capture variability in three spatial dimensions. The pattern of values across sulci and hemispheres, however, is consistent across surface and volumetric probability maps. For all sulci, hemispheric differences were observed with regard to the location in volume space (Fig. 3b). All sulcal segments in the left hemisphere are located consistently further posterior compared to the segments on the right hemisphere, which is in line with the Yakovlevian anticlockwise torque of the two hemispheres (see Table 2 for coordinates of the voxel of maximal probability and for the center of gravity for each label).

**Fig. 3.**
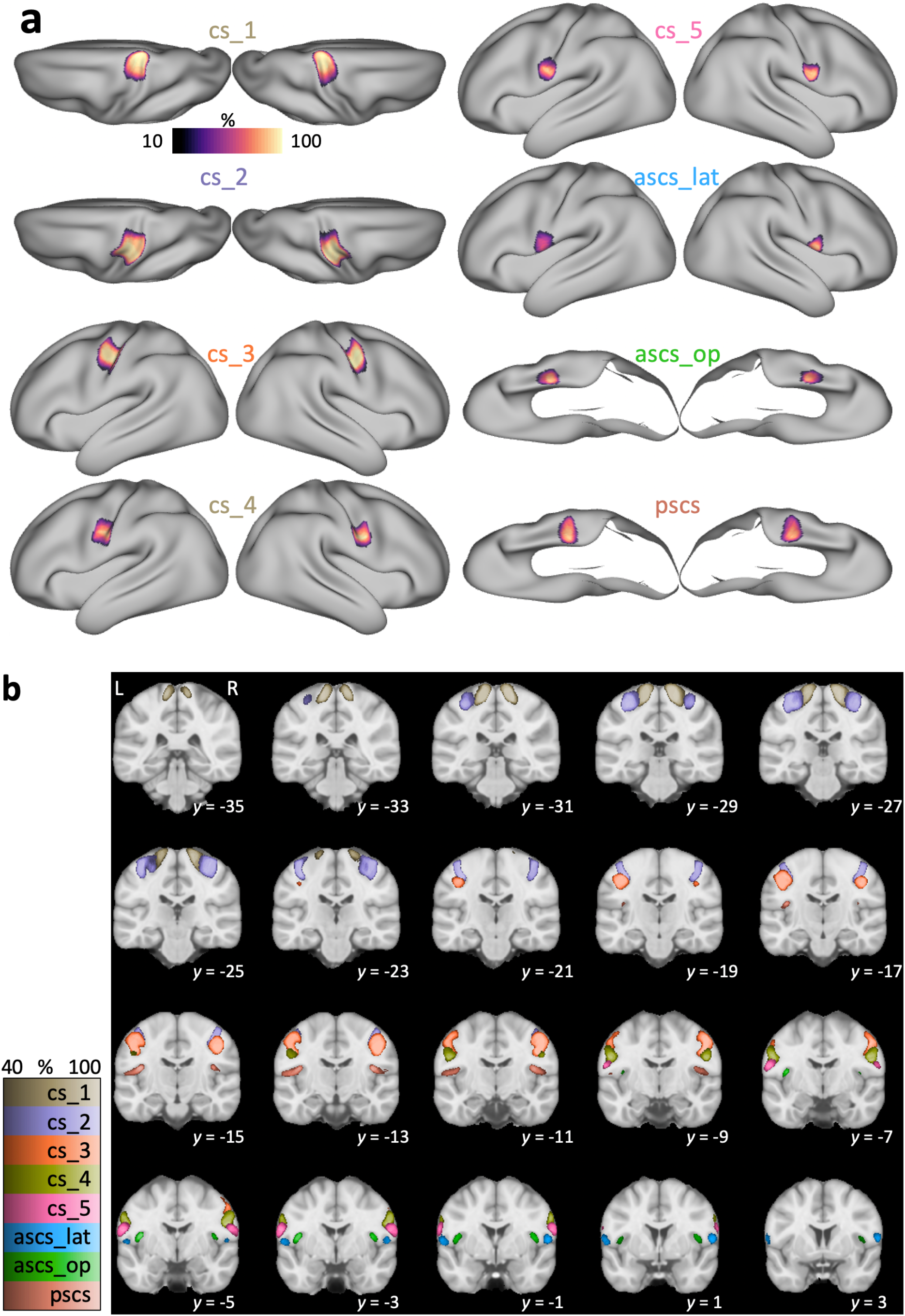
Spatial probability maps. **a** Surface probability maps of central and subcentral sulcal segments shown on an inflated average surface (*n* = 50). Labels for *cs_1* and *cs_2* are shown from a dorsal perspective. Labels for *ascs_op* and *pscs* are shown from a ventral perspective with the temporal lobe removed for better visibility of the opercular cortex. Intensity values show the extent of overlap with 100 % indicating an overlap in all participants. **b** Volumetric probability maps overlaid onto the standard MNI template (*n* = 50). L: left hemisphere shown on left side of the image. Intensity values show the extent of overlap across participants. Color maps for the different sulci have been matched in luminance so that brighter colors indicate higher overlap

**Table 2.** Coordinates of spatial probability maps. For each sulcal segment in left and right hemispheres, we provide the MNI stereotaxic space coordinates (*x, y, z*) for the voxel of maximal overlap and the center of gravity in the volumetric probability maps.

| sulcus | left hemisphere |  |  |  |  |  | right hemisphere |  |  |  |  |  |
| --- | --- | --- | --- | --- | --- | --- | --- | --- | --- | --- | --- | --- |
|  | maximum |  |  | center of gravity |  |  | maximum |  |  | center of gravity |  |  |
|  | X | Y | z | X | y | Z | x | y | z | x | y | Z |
| *cs_1* | -14 | -30 | 69 | -17 | -30 | -69 | 16 | -28 | 68 | 18 | -28 | 69 |
| *cs_2* | -36 | -26 | 58 | -36 | -24 | 59 | 38 | -23 | 60 | 36 | -21 | 58 |
| *cs_3* | -38 | -18 | 42 | -47 | -15 | 46 | 44 | -14 | 40 | 48 | -12 | 46 |
| *cs_4* | -54 | -8 | 30 | -55 | -7 | 32 | 58 | -4 | 32 | 56 | -5 | 32 |
| *cs_5* | -58 | -6 | 22 | -60 | -4 | 22 | 62 | -2 | 21 | 62 | -2 | 22 |
| *ascs_lat* | -60 | 2 | 12 | -58 | 0 | 12 | 58 | 1 | 10 | 58 | 2 | 12 |
| *ascs_op* | -41 | -4 | 14 | -44 | -3 | 10 | 42 | -3 | 14 | 44 | -2 | 11 |
| *pscs* | -39 | -16 | 18 | -48 | -14 | 15 | 41 | -14 | 18 | 48 | -13 | 16 |

The first three segments of the central sulcus (*cs_1* to *cs_3*) show high inter-individual spatial consistency, which is characterized by high values, i.e. overlap, in the probability maps. Consistency is lower for *cs_4* and *cs_5*. Maximal values in the probability map for quantitative comparisons are provided in Fig. 4. Consistency is also low for the subcentral sulci (*ascs_lat*, *ascs_op*, *pscs*) with lowest consistency for *ascs_lat*, especially in the left hemisphere. The hemispheric effect for the *ascs_lat* is evident in both surface and volumetric probability maps. It was observed that the *ascs_lat* is more variable in the left hemisphere (see for example Fig. 5, slice *y* = 1). The low consistency in the *ascs_lat* is in line with the variable morphological types that were described above. The two scentral sulci, *ascs_op* and *pscs*, are less consistent than the central sulcus segments, but more consistent than the *ascs_lat*.

**Fig. 4.**
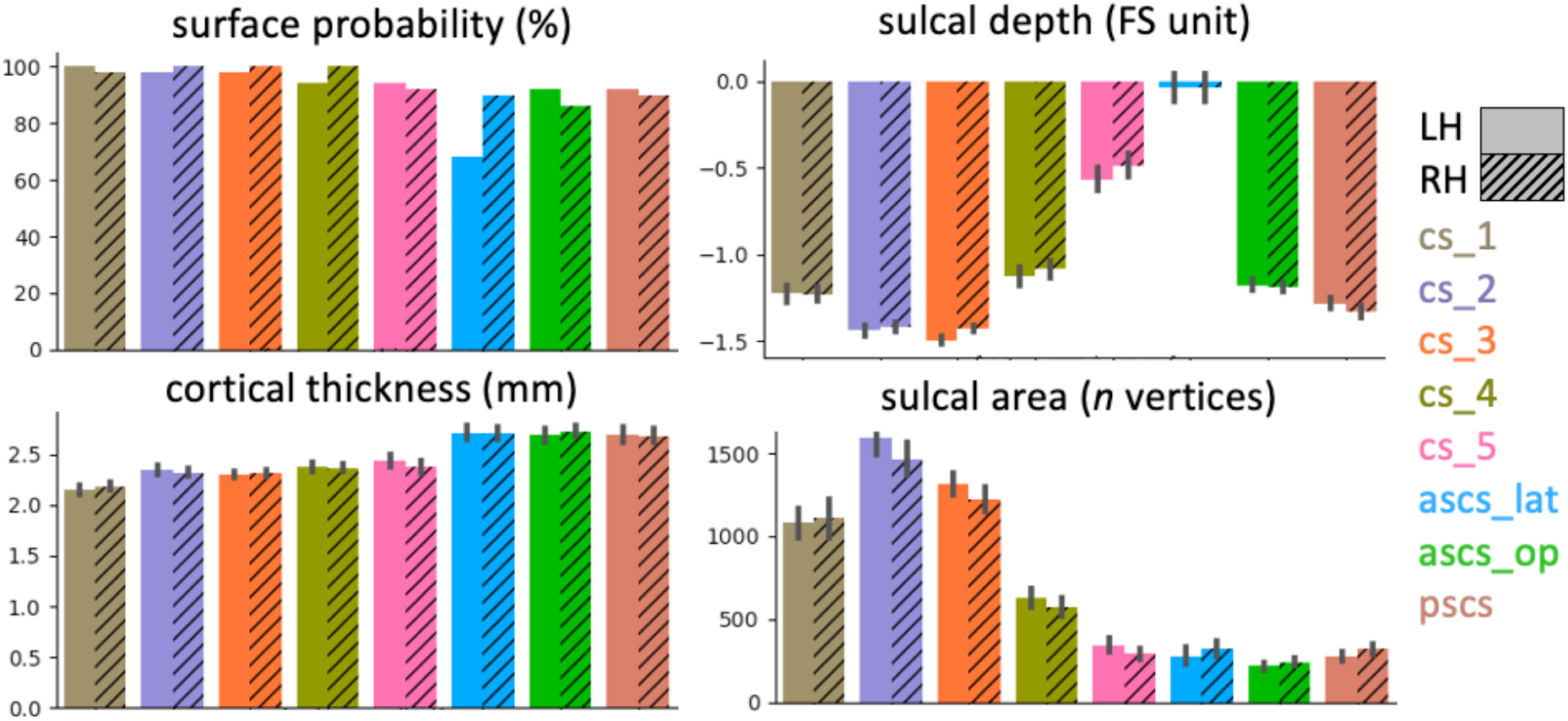
Anatomical measures for sulcal segments. Surface probability quantifies the absolute value of maximal percentage for the overlap of sulcal labels as shown in Fig. 3a. A low probability indicates high inter-individual variability. A more negative sulcal depth indicates a deeper sulcus. Bars for sulcal depth, cortical thickness and sulcal area show mean values ± 95 % confidence intervals (*n* = 50 for each hemisphere)

**Fig. 5.**
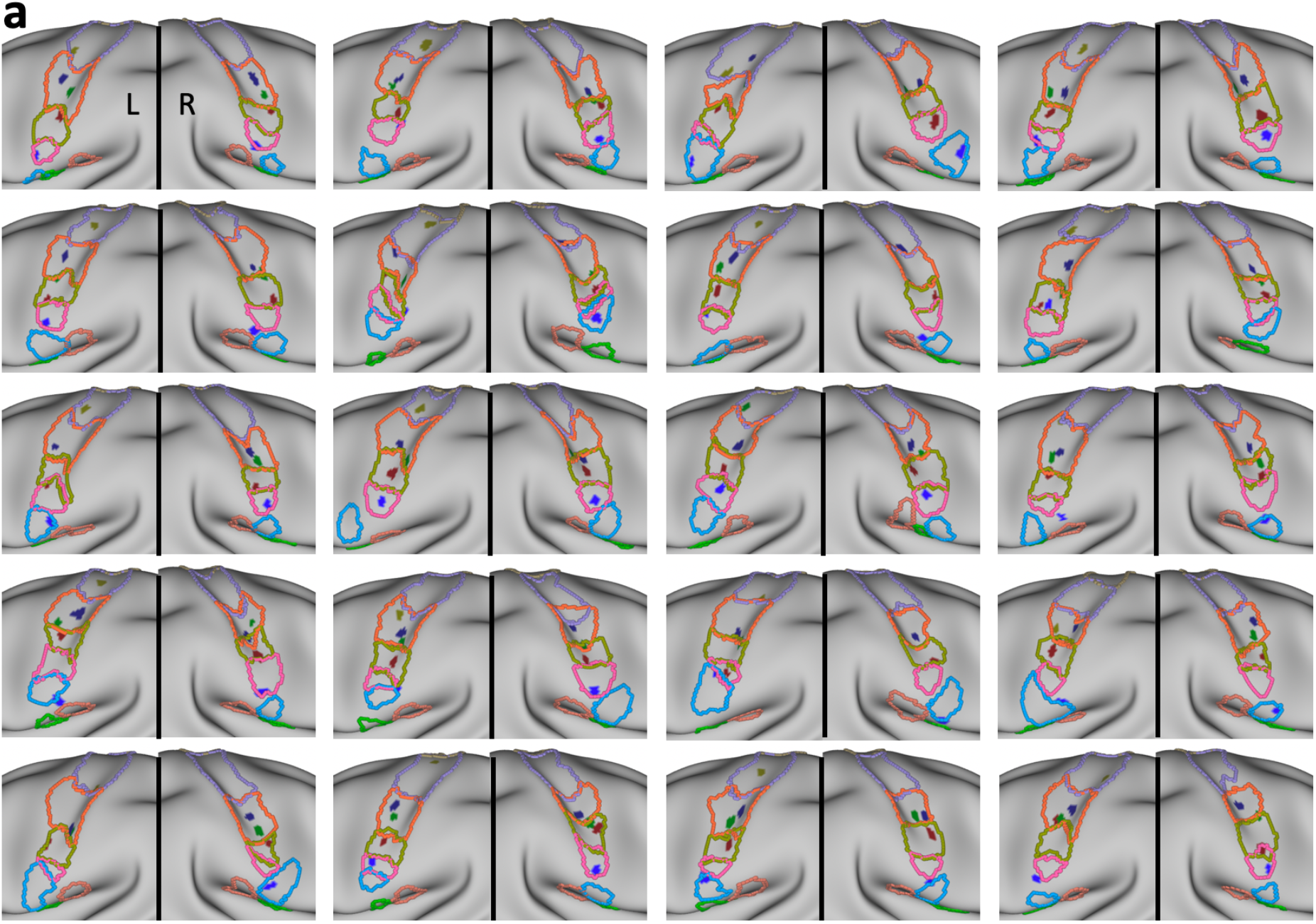
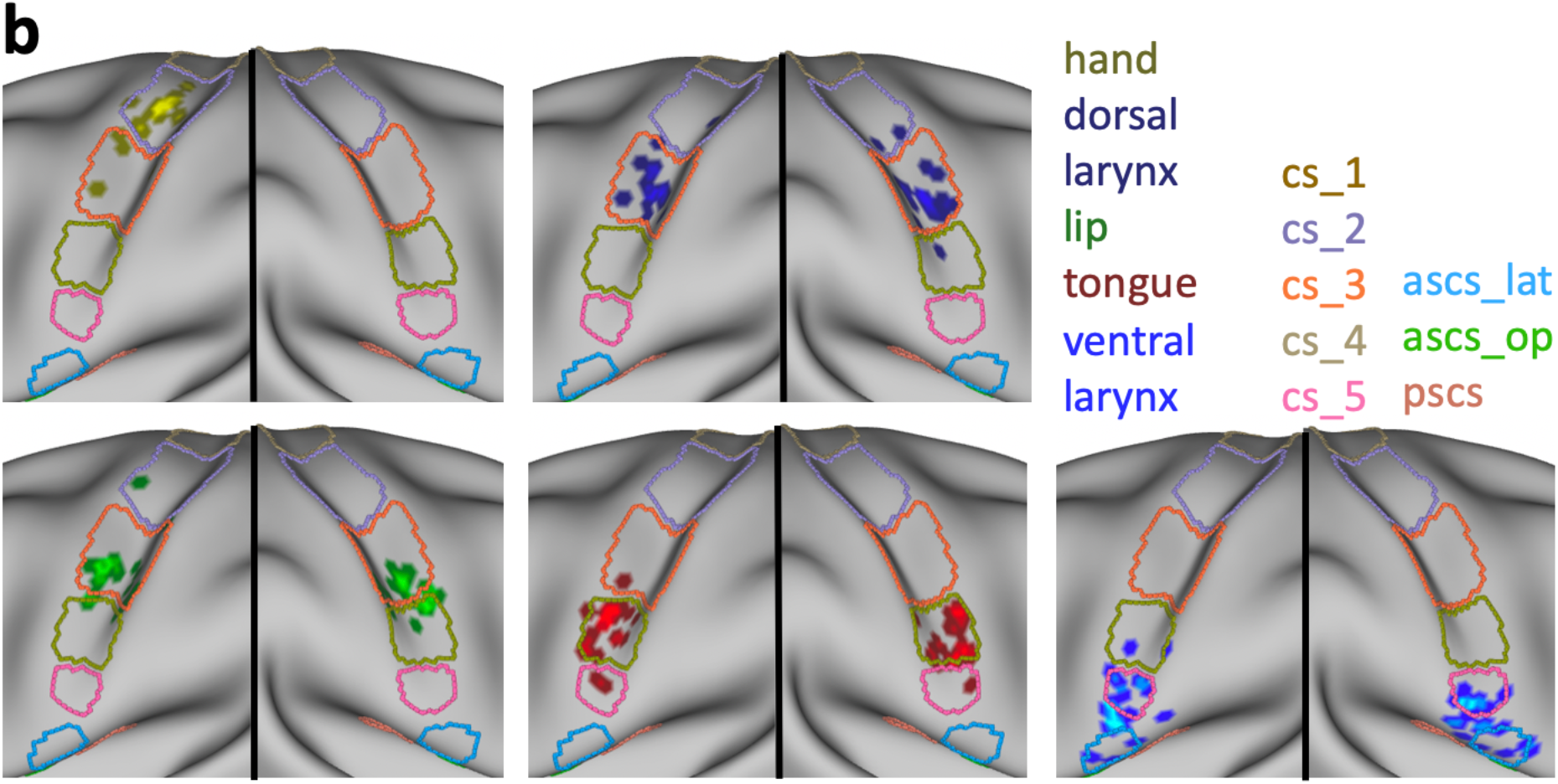
Structure-to-function relationships. **a** Individual sulcal labels and activation maxima for participants in cases where both anatomical and functional data were available (*n* = 20). Only the central strip of the brain is shown. For color legend see b. **b** Location of individual activation peaks with respect to the sulcal maps after applying the sulcal registration (*ascs_op* is not visible). The sulcal outlines show the averaged, normalized and thresholded (> 0.4) sulcal labels after registration. Brighter colors indicate overlapping peaks

### Anatomical Characterization of Sulcal Segments

In addition to probability maps, we characterized the sulcal segments based on sulcal depth, cortical thickness, and sulcal area of the surface label (Fig. 4). It can be observed, that the first three central sulcus segments (*cs_1* to *cs_3*) are deeper than *cs_4* and *cs_5*. Note that the sulcal depth measures are expressed in normalized FreeSurfer units, rather than in real-world values. The *ascs_lat* is notably shallower than the other segments. Cortical thickness increases from dorsal to ventral and is highest in the two subcentral sulcal segments (*ascs_op, pscs*). The largest sulcal segment, as indicated by sulcal area, is *cs_2*. It can be observed that *cs_2* is larger in the left hemisphere, which is consistent with previous reports of a structural asymmetry in the size of the ‘hand knob’ (Yousry et al. 1997; Germann et al. 2019). *Cs_4* and *cs_5* are notably smaller than the other central sulcus segments. None of the three measures described show a hemispheric difference for *ascs_lat*, unlike the surface probability maps, which demonstrate greater variability, i.e. less overlap, in the left hemisphere. Given that handedness of the Human Connectome Project (HCP) participants was not accessed, we did not assess how this might have affected the measures described above. Regardless of handedness, however, it must be assumed that the majority of the 50 participants would be left lateralized for language. In the supplementary material, we report the anatomical measures depending on the morphological type of the subcentral region (Supplementary Material, Fig. 8). Although minor deviations from the patterns described in Fig. 4 can be observed, overall, the anatomical characteristics for the sulcal segments are similar across all morphological types.

### Sulcal Registration and Structure-to-Function Relationship

All individual sulcal labels and functional activation peaks are shown in Fig. 5a. To visualize the spatial relationships at the group level, we first registered all individual surfaces based on the binary sulcal labels using MSM. After registering the sulcal labels, we averaged all transformed sulcal labels and thresholded (> 0.4) the average labels to obtain the outlines of registered sulcal segments. Fig. 5b shows the outlines of the registered sulci, which demonstrates an orderly dorsal-ventral configuration of the sulcal segments.

Next, we applied the registration that was used to transform individual surface labels to the group space to the functional activation peaks of different effectors in individual participants. Fig. 5b shows the location of the resampled activation maxima in relation to the outlines of the registered segments. Given that the same registration was applied to individual sulcal labels and the functional peaks, the location of resampled peaks with respect to the registered sulci reflects the original spatial relationship on the native surfaces of the individuals. The location of the peaks is therefore similar, but not identical to the location obtained from a group-level FreeSurfer-based registration (as shown in Figure 4A in Eichert et al. 2020a).

Overall, a strong correspondence between sulcal labels and functional peaks was observed. The observed relationships between labels and peaks are reported in Table 3. The majority of activation peaks from the hand localizer fall inside the center of *cs_2*. The dorsal peaks for larynx activity are found on the anterior bank of *cs_3*. Peaks from the lip localizer also fall within *cs_3* and overlap with the location of the dorsal larynx peaks. Those for the lip, however, are reliably located at the more ventral extent of *cs_3* whereas those for the dorsal larynx peaks span the segment. Activation peaks from the tongue localizer fall inside *cs_4* with high consistency. The ventral peaks for larynx activity are associated with *cs_5* and *ascs_lat* or with the gyrus in between, but the relationship is less consistent than for the other functional peaks. The sub-threshold maxima were similar in terms of their location to the other maxima.

**Table 3.** Structure-to-function relationships. The numbers show the observed relationships between functional activation peaks (*n* = 20) and sulcal label underlying the peak depending on the hemisphere.

|  | LH |  |  |  |  | RH |  |  |  |
| --- | --- | --- | --- | --- | --- | --- | --- | --- | --- |
|  | Hand | dorsal larynx | Lip | Tongue | ventral larynx | dorsal larynx | Lip | tongue | ventral larynx |
| *cs_1* | 0 | 0 | 0 | 0 | 0 | 0 | 0 | 0 | 0 |
| *cs_2* | 16 | 1 | 1 | 2 | 0 | 1 | 0 | 0 | 0 |
| *cs_3* | 4 | 19 | 19 | 1 | 0 | 18 | 15 | 0 | 0 |
| *cs_4* | 0 | 0 | 0 | 17 | 2 | 1 | 5 | 19 | 0 |
| *cs_5* | 0 | 0 | 0 | 0 | 12 | 0 | 0 | 1 | 12 |
| *ascs_lat* | 0 | 0 | 0 | 0 | 6 | 0 | 0 | 0 | 8 |

The relationship between sulcal label and effector is very similar across hemispheres (Table 3). The only exception is the lip cluster in the right hemisphere, where several peaks are located in *cs_4* rather than in *cs_3*. The group-level representation (Fig. 5b), however, demonstrates that this difference is driven by minor variations in the boundary between *cs_4* and *cs_3*, rather than being a systematic hemispheric difference in the cortical location of the activity. The sulcal label underlying the ventral larynx peak was the same in both hemispheres for the majority of participants (*n* > 12). Furthermore, we examined the relationship between morphological type of the subcentral region and the location of the ventral larynx peak. We observed no systematic relationship, i.e. all combinations of type and sulcal label are present. Given the low count number for each combination, a robust quantification of this observation is not feasible.

To characterize the effect of the sulcal registration on the spatial variability of functional peaks, we computed the median distance across all peaks for each effector. We compared this measure of spatial spread with that obtained from the FreeSurfer registration and a registration based on sulcal depth maps. Overall, the three registrations have a similar effect on the spatial variability (Fig. 6a). Some differences can be observed across effectors, but given the sample size none of the effects are significant. For the lip cluster in the left hemisphere, spatial spread is increased for the registration based on sulcal labels. Spatial variability of the ventral larynx cluster in the left hemisphere was reduced the most by using the sulcal-segment registration, but the effect is small and not statistically significant. Fig. 6b shows the spatial distribution of the fMRI peaks for the ventral larynx representation after a FreeSurfer-based registration and after registration based on sulcal segments. The peaks in the left hemisphere appear more focal when sulcal segments are driving the registration, which is in line with a reduction of the median distance. In the right ventral larynx cluster, however, the median distance was overall increased and the overlap of fMRI peaks decreased. The spatial map, however demonstrates that the registration brings several outlier peaks closer to the remaining cluster.

**Fig. 6.**
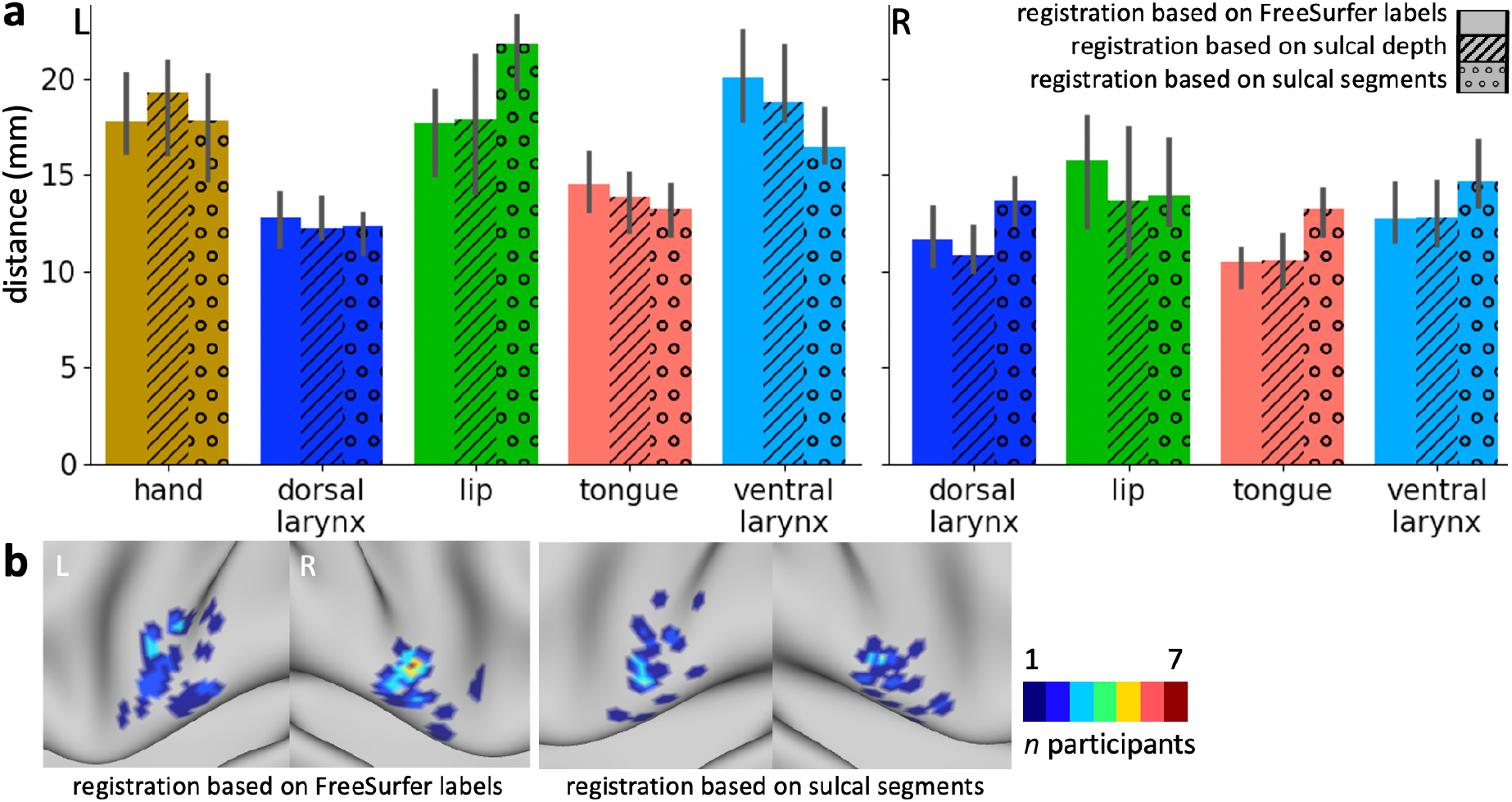
Effect of anatomical registrations on functional variability. **a** Distance of task activation peaks after registration based on FreeSurfer labels (blank bars), after registration based on whole-brain sulcal depth maps (striped bars) and after registration based on manually drawn sulcal segment labels (dotted bars). The median distance ± 95% confidence interval is shown across all peaks for each task cluster (*n* = 20; 190 pairs of points for each cluster). **b** Overlap of fMRI peaks of the ventral larynx representation after FreeSurfer registration and after registration based on sulcal segments.

## Discussion

We examined the morphological patterns and the inter-individual variability of the sulci in the central and subcentral region of the cerebral cortex in the human brain and their relation to activity evoked by movement. A robust relationship between morphological features of the central and subcentral sulcal segments and movement of different effectors at the individual participant level was demonstrated. An inter-individual registration based on the individually drawn sulcal segments demonstrated that morphological variability can explain functional variability, in particular in the subcentral region, where morphology is complex.

### Morphology of the Subcentral Cortex and the Central Operculum

The present study provided a systematic examination of the sulcal patterns of the subcentral and adjacent opercular cortical region in the human brain, including inter-individual variability of the sulcal morphology. In this region, one encounters an anterior and a posterior subcentral sulcus (*ascs*, *pscs*) (Petrides, 2012, 2019). We showed that, in the majority of participants, the *ascs* consists of a distinct lateral (*ascs_lat*) and a distinct opercular segment (*ascs_op*).

The morphological patterns of the subcentral region were classified into five distinct types, based on the configuration of the *ascs_lat*. In addition to the canonical configuration (Type 1, Fig. 1b), the *ascs_lat* often merged with the central sulcus, so that the central sulcus complex extended further ventral towards the lateral fissure (Type 2). In Type 3, the position of the *ascs_lat* is notably more opercular and it merges with the *ascs_op*, so that the *ascs_lat* and *ascs_op* form one continuous sulcus. In Type 4, the *ascs_lat* merges with the inferior precentral sulcus (*iprs*). Type 5 is characterized by a more anterior position of the *ascs_lat*, compared with its position in the other types. The variability in the configuration of the lateral and opercular segments explains why the *ascs* appears as one continuous sulcus in a volumetric average of structural brain images in MNI stereotaxic space (Petrides, 2019).

### Sulcal Labeling in Surface Space

We aimed to reproduce the segmentation of the central sulcus, as described in a volumetric labeling study (Germann et al. 2020). We labeled sulcal segments directly on the participant’s native cortical surface, rather than in volume space as traditionally done in several MRI-based labeling studies (Germann et al. 2005, 2020; Zlatkina and Petrides 2010; Amiez et al. 2013; Sprung-Much and Petrides 2018, 2020). We demonstrated the advantages of working in surface space as this can explicitly take into account the geometry of the cortical convolutions.

The anatomical landmarks critical for labeling are conveniently visualized in surface space and, therefore, are more easily reproducible (Fig. 1b). We characterized the sulcal segments using several surface-based anatomical features. Furthermore, quantifications of geometrical and topological properties of interest for sulcus morphometry are readily available for surface labels (Mangin et al. 2004). Also, using surface labels allowed us to perform a surface-based registration, where distances are represented as geodesic distances along the cortex rather than as Euclidean distances between voxels (Fischl et al. 1999b; Klein et al. 2010). In the current study, the labels were drawn manually, but the development of automatic labeling algorithms for sulcal segments is subject of ongoing research (Clouchoux et al. 2006; Takerkart et al. 2015; Machlouzarides-Shalit et al. 2020; Borne et al. 2020).

### Spatial Probability Maps

We visualized the morphological variability and spatial extent of the five segments of the central sulcus (*cs_1* to *cs_5*) and the subcentral sulci in surface and volumetric probability maps. The first three central sulcus segments show the highest spatial consistency across participants. Consistency decreases for the fourth and even further for the most ventral (fifth) segment of the central sulcus, which is in line with previous observations (Germann et al. 2020). The decrease in probability values for *cs_4* and *cs_5* can also be attributed to their smaller spatial extent compared with the sulcal labels for *cs_1* to *cs_3*. Consistency for the subcentral sulci *ascs_lat*, *ascs_op* and *pscs* is also notably reduced, which is in line with the inter-participant variability in the morphological patterns as described above.

### Hemispheric Differences

The classification of morphological types and the spatial probability maps demonstrated hemispheric differences in the subcentral cortical region, namely that the position of the *ascs_lat* along the rostro-caudal axis differs between hemispheres, but not its dorso-ventral position. The *ascs_lat* in the left hemisphere tends to be located further posteriorly, which frequently results in a merge with the central sulcus, while the *ascs_lat* in the right hemisphere is located more anteriorly. The present quantifications showed that the *ascs_lat* is more variable in the left hemisphere, although several anatomical measures of the *ascs_lat* are not different across hemispheres, such as the sulcal depth and sulcal area.

The hemispheric difference in variability can be interpreted in relation to sulcal variability of the ventro-lateral cortex (Germann et al. 2005; Sprung-Much and Petrides 2018, 2020). Language processing is lateralized most frequently to the left hemisphere (language dominant hemisphere) and the role of the left inferior frontal cortex in language function is widely established (Broca 1861; Price 2000). Functional language lateralization is associated with structural asymmetries (Foundas et al. 1996; Josse and Tzourio-Mazoyer 2004) and increased regional variability (Croxson et al. 2018), but the structure-to-function relationships remain controversial (Dorsaint-Pierre et al. 2006; Sprung-Much and Petrides 2018). It is possible that that increased surface area in the left inferior frontal cortex affects the neighboring sulci so that the *ascs_lat* is ‘pushed’ to a relatively more posterior location in the left hemisphere. Despite the hemispheric differences in morphology, the structure-to-function relationships for basic movements of different effectors described below, did not exhibit notable differences between hemispheres.

Several previous studies indicated that structural asymmetries in the human motor cortex are related to handedness and gender (Amunts et al. 2000; Mangin et al. 2004; Sun et al. 2012; Germann et al. 2019). These effects are not limited to the hand knob but affect the shape of the entire central sulcus. Future investigations could test for a systematic relationship between handedness and the morphological variations described here for the subcentral cortical region. Linking functional and morphological variability to behavioral measures would improve our understanding of hemispheric specializations underlying human language and handedness.

### Functional Activation Peaks and Morphology

Here, we demonstrated a tight link between individual morphological features of the cortex and activation peaks for the different effectors examined. We replicated the relationships between segments of the central sulcus and functional localizers for the hand, lip, tongue, and larynx reported in Germann et al. (2020). We identified two larynx peaks, consistent with more recent studies suggesting that two separate regions in the central region correlate with larynx activity (Belyk and Brown 2017; Jarvis 2019; Eichert et al. 2020a). Germann et al. (2020) however, found only a ventral larynx peak. The difference across studies might be because our functional localizer differed substantially from the one used by Germann and colleagues In the latter study, ‘humming’ was used as task instruction to localize larynx activity. Germann et al. found the ventral larynx peak to be associated with *cs_5*, and did not identify *ascs_lat*. We showed that it was also localized in the *ascs_lat* in a large proportion of participants (14 out of 40 hemispheres). This difference might be due to the significantly larger sample size in the current study that allowed a more robust assessment of the relationships. A systematic location of the dorsal larynx peak relative to morphological features has been predicted before (Sun et al. 2016), but this is the first study that describes consistent structure-function relationships for both the dorsal and the ventral larynx representations and how they relate to the representations of the other articulators.

It should be noted that the functional contribution of the two larynx representations in motor control during vocalization is still debated (Simonyan 2014; Belyk and Brown 2017). Quantifications of cortical microstructure, such as myelin content, indicate that the ventral larynx region is not located in primary motor cortex (Eichert et al. 2020a). The focus of the current study, however, is not the interpretation of the functional activation peaks *per se*, but rather the assess of the relationship between their location and the underlying morphology. The individual variability in the cortical location of the ventral larynx representation could be one factor in explaining inconsistent reports in the literature.

The morphological type was not predictive of the sulcal label underlying the ventral larynx peak and vice versa. At the given sample size of 20 participants, the total counts for each combination of morphological type and sulcal label is low and, therefore, it is not practical to quantify the robustness of this effect. The current dataset, however, suggests that the sulcal label predicts the location of the maximal functional activation, regardless of the morphological configuration formed by the sulci. Future research could elaborate on the functional significance of the morphological types (Sun et al. 2016). For example, using an fMRI-paradigm that activates the opercular region more directly might reveal effects on the size of the activated area or the activation strength.

### Relationship between Variability in Structure and Function

We observed a robust relationship between sulcal landmarks and the location of functional activation peaks for speech-related effectors. The somatomotor system, however, is only one example system, where a link between sulcal morphology and function at the individual participant level has been established (Boling et al. 1999; Coulon et al. 2011; Zlatkina et al. 2016; Germann et al. 2020). Detailed anatomical and functional studies have also revealed such relationships for other parts of the cortex (e.g., Amiez et al. 2006; Bodin et al. 2018).

A registration based on individually drawn sulci reduced the spatial spread of task peaks for the left ventral larynx representation. This indicates an improved registration in this part of the brain, when compared to a registration based on sulcal depth and based on FreeSurfer. The effect, however, was small and not statistically significant, and for other effectors the registration even increased the spatial spread. Reducing the whole-brain statistical task activation maps to a peak might have obscured greater benefits of a registration based on sulcal segments.

In many contexts, such as brain development or cross-species comparisons, the manual identification of sulcal landmarks is vital to constrain the registration algorithm when automatic labeling is inaccurate or not feasible (Van Essen and Dierker 2007; Lebenberg et al. 2018; Eichert et al. 2020b). Here we show that manual identification of sulcal landmarks is also critical for registrations in the human brain, particularly in brain regions that are highly variable and have low gyrification. The ventral larynx peaks were not consistently co-localized with any specific sulcal segment, but were located in both *cs_5* and *ascs_lat*. The anatomical information of the labels, however, was still helpful for the alignment within the subcentral opercular region. The current study emphasizes the importance of detailed anatomical investigations of sulcal segments and tertiary sulci in understanding the structure-to-function relationships in the cortex (Weiner et al. 2018; Miller et al. 2020).

In the current study, we examined gross anatomical features of the brain surface, an approach that is directly applicable to neuronavigation and neurosurgical planning. Recent advances in neuroimaging, however, allow us to acquire multiple image modalities in the same participants, some of which correlate with the underlying cellular microstructure (Fischl and Dale 2000; Zhang et al. 2012; Weiskopf et al. 2013). Although the histological analysis of post-mortem brains remains the gold standard to define anatomical parcels of the brain, neuroimaging has become a versatile tool for parcellating the cortex and to establish relationships between different modalities (Glasser et al. 2016; Jakobsen et al. 2018; Smith et al. 2019; Eichert et al. 2020b). Future work will show how detailed morphological labeling and segmentation studies can benefit from other neuroimaging modalities.

In summary, the current study demonstrated a robust relationship between morphological features of the central and subcentral sulcal segments and movement of different effectors at the individual participant level. We described the morphological patterns of the sulci in the subcentral and central opercular cortical region, their inter-individual variability and several anatomical properties. Laryngeal activity during vocalization activates the subcentral region, which we found to be highly variable across participants. This variability could explain inconsistencies in previous reports about a ventral larynx representation.

## Supplementary Material

### Anatomical Characterization of Sulcal Segments based on Morphological Types

**Fig. 8.**
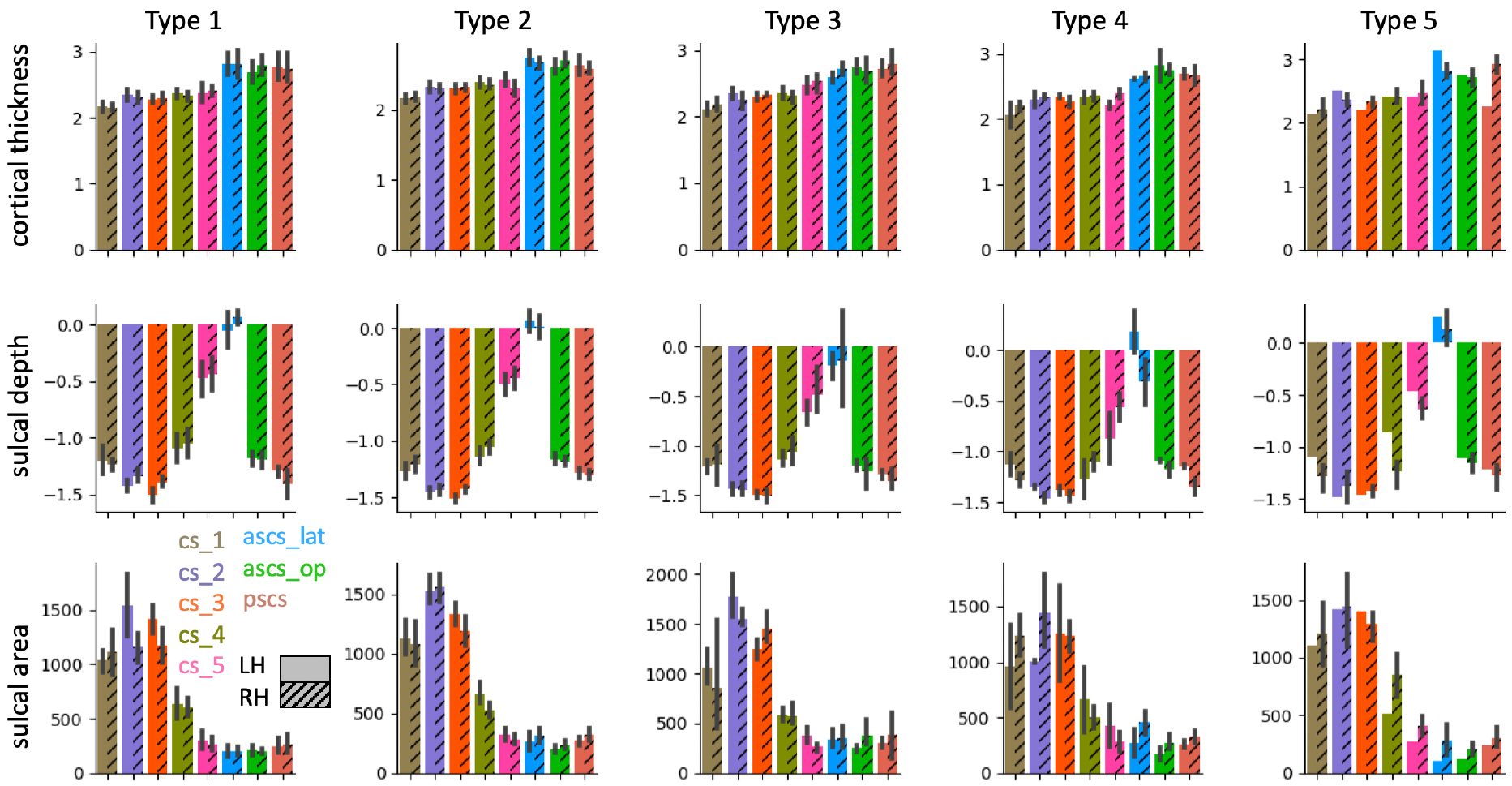
Anatomical measures for sulcal segments in different morphological types. Morphological types refer to the patterns in the subcentral region as shown in Fig. 2. For more details on the figure description see Fig. 4. Sample sizes underlying each type and hemisphere are provided in Table 1

### Regions-of-Interest to derive Maxima during Task Activation

The definition of regions of interest is described, in detail, in Eichert et al. (2020a) and is repeated here in a modified version. The central sulcus ROI used for the hand, lip and tongue was defined using FreeSurfer’s automatic volumetric labeling based on the Destrieux Atlas.

For the larynx, we identified two activation maxima in separate ROIs: One for the dorsal and one for the ventral larynx representation. The dorsal larynx ROI was a portion of the same central sulcus ROI used above from *z*-coordinates in MNI space of 50 - 30. The limits were determined empirically, so that the dorsal larynx ROI did not capture the ventral larynx representation or an unrelated supra-dorsal activation in the trunk area, which was observed in some individuals (Foerster 1931).

The ventral larynx representation lay outside the central sulcus and was located ventrally in the subcentral part of cortex. Because of the high inter-individual morphological variability in this region, the ventral larynx ROI was derived manually based on individual anatomy in surface space. A liberal surface ROI was drawn on each individual’s mid-thickness surface covering the ventral part of the central sulcus and adjacent gyri. Anteriorly, the ROI was delineated by the inferior portion of the precentral sulcus and, posteriorly, the ROI spanned the postcentral gyrus. If present, the lateral portion of the anterior sulcus in the subcentral gyrus was included within the ROI. The dorsal limit of the ROI was defined by a horizontal plane across the gyrus at the level of the usual location of the posterior ramus of the inferior precentral sulcus. The ventral larynx surface ROI was converted into a volumetric ROI covering the underlying cortical ribbon using wb_command. We checked that the ventral larynx ROI did not overlap with subjacent auditory cortex in the temporal lobe or inferior frontal cortex.

In some participants, the main contrast for vocalization in the syllable production task had additional activity related to articulation of the tongue. To remove this additional activity, we transformed the coordinates for each individual’s maximal voxel from the tongue contrast (from the basic localizer task) to the functional space of the syllable production task (task 1) using rigid-body transformation and then derived a spherical ROI (7 voxels in diameter) around it. This sphere was used to mask the *z*-statistic image of the main contrast for vocalization prior to localizing the maxima for laryngeal activity in the dorsal and ventral ROIs described above.

### Animation of one Individual’s Sulcal Segments

#### Online Resource 1

The animation shows the 3D rendering of one individual’s sulcal segments in native volume space, i.e. the same participant as in Fig. 1 using the same color legend.

## Declarations

### Compliance with Ethical Standards

HCP participants: The Ethical standards are fully described in the core HCP literature referenced in the main body. The remaining participants were part of a study that had been approved by the Central University Research Ethics Committee of the University of Oxford (CUREC, R55787/RE001) in accordance with the regulatory standards of the Code of Ethics of the World Medical Association (Declaration of Helsinki). All participants gave informed consent for their participation.

### Availability of data and material

Anatomical raw data of the participants that provided structural and functional data are publicly available at OpenNeuro under the accession code ds002634. The minimally pre-processed data of the HCP participants are openly available for download at https://db.humanconnectome.org. Upon acceptance of the manuscript, derived data supporting the findings of this study, including the individual surface renderings and sulcal labels, will be made available from the Wellcome Centre for Integrative Neuroimaging’s GitLab at git.fmrib.ox.ac.uk/neichert/project_variability.

### Code availability

Upon acceptance of the manuscript, processing code will be made available from the Wellcome Centre for Integrative Neuroimaging’s GitLab at git.fmrib.ox.ac.uk/neichert/project_variability. Parts of the processing code is already available at git.fmrib.ox.ac.uk/neichert/project_larynx. FSL tools, including MSM, are available from fsl.fmrib.ox.ac.uk. Connectome Workbench is available at www.humanconnectome.org/software/connectome-workbench.html.

### Author Contributions

Conception and design: NE, MP. Acquisition of data: NE, KEW. Analysis and interpretation of data: NE, KEW, RBM, MP. Original draft: NE, MP. Revision of the article: NE, KEW, RBM, MP. Contribution of analytic tools: NE, RBM.

